# Adaptation to soil type contributes little to local adaptation in an Italian and a Swedish population of *Arabidopsis thaliana* growing on contrasting soils

**DOI:** 10.1101/2024.04.24.590956

**Authors:** Thomas James Ellis, Jon Ågren

## Abstract

Natural populations are subject to selection caused by a range of biotic and abiotic factors in their native habitats. Identifying these agents of selection and quantifying their effects is key to understanding how populations adapt to local conditions. We performed a fully factorial reciprocal transplant experiment using locally adapted accessions of *Arabidopsis thaliana* at their native sites to distinguish the contributions of adaptation to soil type and climate. Overall adaptive differentiation was strong at both sites. However, we found only very small differences in the strength of selection on local and non-local soil, and adaptation to soil type at most constituted only a few percent of overall adaptive differentiation. These results indicate that adaptation to local climatic conditions rather than soil type is the primary driver of adaptive differentiation between these ecotypes.

## Introduction

Divergent selection due to differences in biotic, climatic and other physical conditions contributes to the maintenance of adaptive genetic diversity among natural populations and the evolution of local adaptation (1; 2). Reciprocal transplant experiments have demonstrated local adaptation across several spatial scales and in many species (2), but in most cases the agents responsible for divergent selection have not been established. This is problematic as a full understanding of the ecological causes of adaptation requires that both adaptive traits and agents of selection are identified. Among-population correlations among traits and environmental factors can suggest the drivers of divergent selection. However, to determine conclusively agents of selection requires studies that adopt an experimental approach (3–5).

Terrestrial plants derive nutrients required for growth predominantly through interactions with the soil and soil biota. Since the biotic and abiotic characteristics of soil vary immensely, these interactions can select for plant traits that promote increased performance on a specific soil type. Adaptations to edaphic conditions are many, including to pH, salinity, carbonate levels, serpentine soil, and soil microbiota (6–13). If plant populations adapt to the edaphic conditions in their local environment, this will contribute to adaptive differentiation between populations (14). In this way, soil type may act as a selective agent contributing to local adaptation.

In this study, we investigate adaptation to native soils in two locally adapted populations of *Arabidopsis thaliana* in Italy and Sweden. The native sites of these populations differ strongly in climate, and previous work using reciprocal transplant experiments of local ecotypes and recombinant inbred lines derived from them have demonstrated local adaptation to climatic conditions (15–19) The soils at these sites are also very different, with higher iron content, lower pH, and lower availability of magnesium and calcium at the Swedish than at the Italian site (13). Nevertheless, neither field experiments at the native sites in Italy and Sweden using the parental ecotypes grown on soil from both source sites, nor a growth-chamber experiment that disentangled the biotic and abiotic components of these soils found evidence for adaptation of plant ecotypes to their native soils (13; 20). However, these studies were limited by relatively small sample sizes and substantial block effects. Moreover, the field experiment was conducted in a single year, and no plants of the Italian ecotype survived to reproduce at the Swedish site, making it difficult to fully evaluate the effect of soil on adaptive differentiation. As such, it is difficult to conclusively rule out a role for adaptation to soil type in local adaptation between these ecotypes.

We conducted a field experiment in which plant ecotypes and soil were reciprocally transplanted between the native sites of the two *A. thaliana* populations to disentangle the contributions of adaptation to soil type and climate. We employ a design that builds on the weaknesses of previous studies by using a fully randomised design with substantially larger sample sizes than those used in earlier work (13; 20). We tested the prediction that if soil type contributes to local adaptation, the fitness advantage of local plants will be greater when grown on the local soil than on the non-local soil.

## Methods

We compared plant fitness in a fully crossed reciprocal-transplant experiment varying site, soil type and plant ecotype. We grew local accessions of *A. thaliana* at sites in central Italy (Castelnuovo di Porto) and north-central Sweden (Rödåsen) on soils collected at each site. Sites, accessions used, and experimental set-up followed those of (15). Briefly, we collected soil from the two source sites in Spring 2017, and stored this in plastic buckets at 6°C. We germinated seeds of each accession on agar in petri dishes in a growth room at Uppsala University, transported these to field sites and transplanted seedlings to 299-cell plug trays with individual cells of 20 mm x 20 mm x 40 mm (HerkuPlast Kubern GmbH, Ering, Germany). We transplanted a total of 2080 seedlings across the two sites, corresponding to 256 and 264 of the local and non-local ecotype on each soil type at each site in September 2017 in Sweden and November 2017 in Italy. In April Spring 2018 in Italy and June 2018 in Sweden, we recorded survival to reproduction, and fruit number of each reproductive plant. For 707 plants in Italy and 488 plants in Sweden, we estimated seed number per fruit by counting all viable seeds from one mature unopened fruit as described in reference (21). We estimated seed number per planted seedling, a proxy for overall fitness, as the product of fruit number and site-soil-ecotype-mean seed number per fruit.

For each site-soil-ecotype treatment, we estimated selection in terms of calculated mean overall fitness, as well as through the survival and fecundity components of fitness. For each site x soil combination, we quantified selection against the non-local ecotype by calculating the selection coefficient as *s* = 1−(*w*_*local*_*/w*_*non*−*local*_), where *w*_*local*_ and *w*_*non*−*local*_ are the mean absolute fitness values for the local and non-local ecotypes respectively. We calculated this for each fitness component and overall fitness separately. Because the fitness data, in particular seed number per seedling planted, were strongly zero-inflated, we quantified 95% confidence intervals around parameter estimates by drawing 1000 bootstrap samples of the data stratified by site-soil-ecotype treatment, and recalculating parameters for each. We calculated p-values for the (two-sided) null hypothesis that there is no difference in selection between the two soil types by calculating twice the proportion of bootstrap samples for which selection on the soil with the strongest observed selection was greater than selection on the other soil. that were greater on the onelocal soil than the othernon-local soil at each site. We multiplied this proportion by two to make this a two-sided test.

We performed statistical analyses in RStudio 2023.06.1+524 using R 4.2.1 (22; 23).

## Results

Overall adaptive differentiation on local soils was strong at both sites (fig. 1). Local ecotypes showed 4.4- and 6.2-fold higher overall fitness (seed number per planted seedling) on local soils in Italy and Sweden, respectively. Local ecotypes also showed increased fitness through all three components of fitness, although selection through survival was stronger in Sweden, and selection through fecundity components stronger in Italy (fig. 2). The patterns of selection found in this study are consistent with previous studies of the same populations (15; 16).

**Figure 1.**
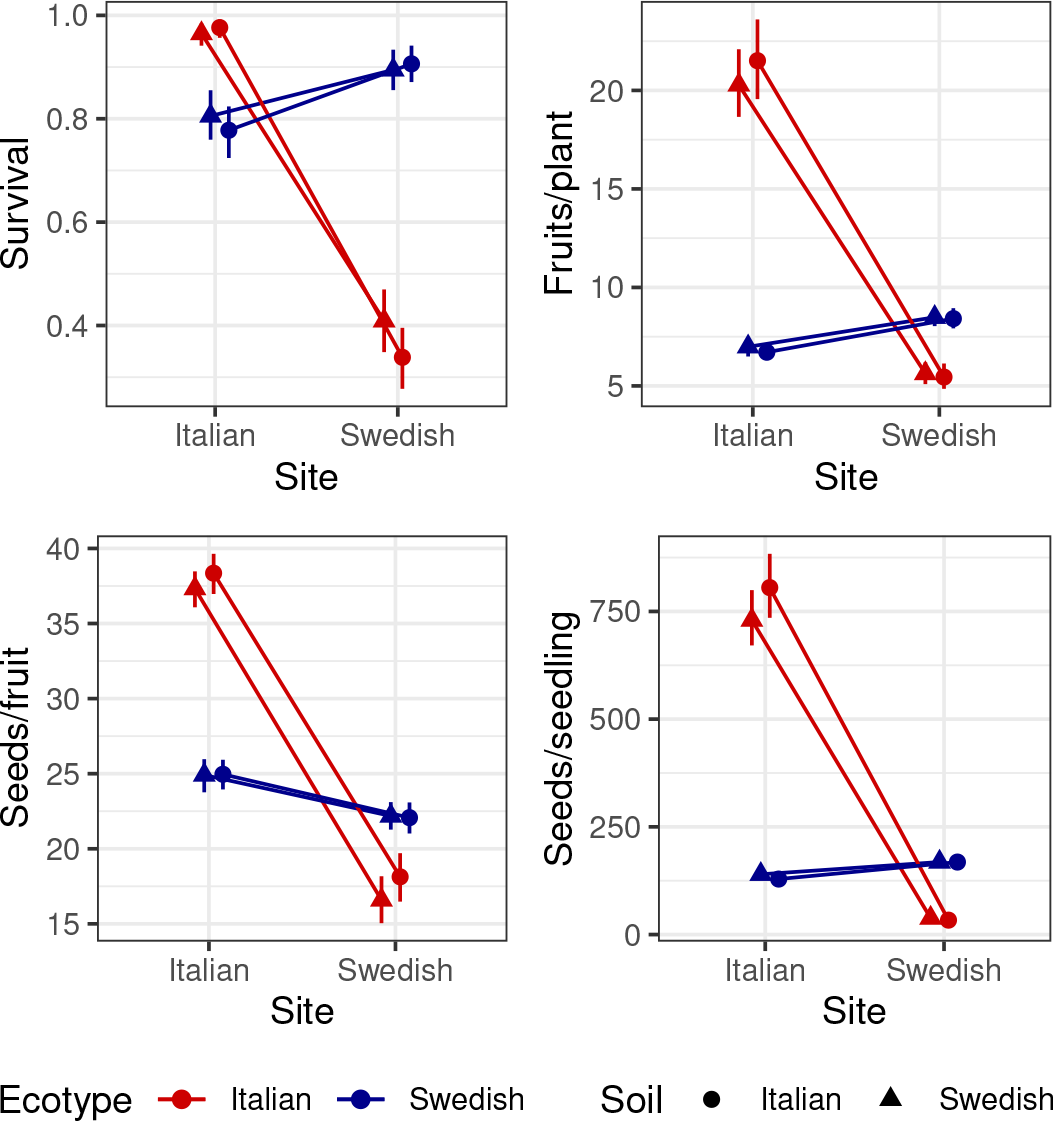
Fitness and its components for Italian and Swedish ecotypes when growing on soils collected at the Italian and Swedish sites. Plots show means and 95% confidence intervals for survival to reproduction, number of fruits per reproductive plant, number of seeds per fruit, and number of seeds per planted seedling.

**Figure 2.**
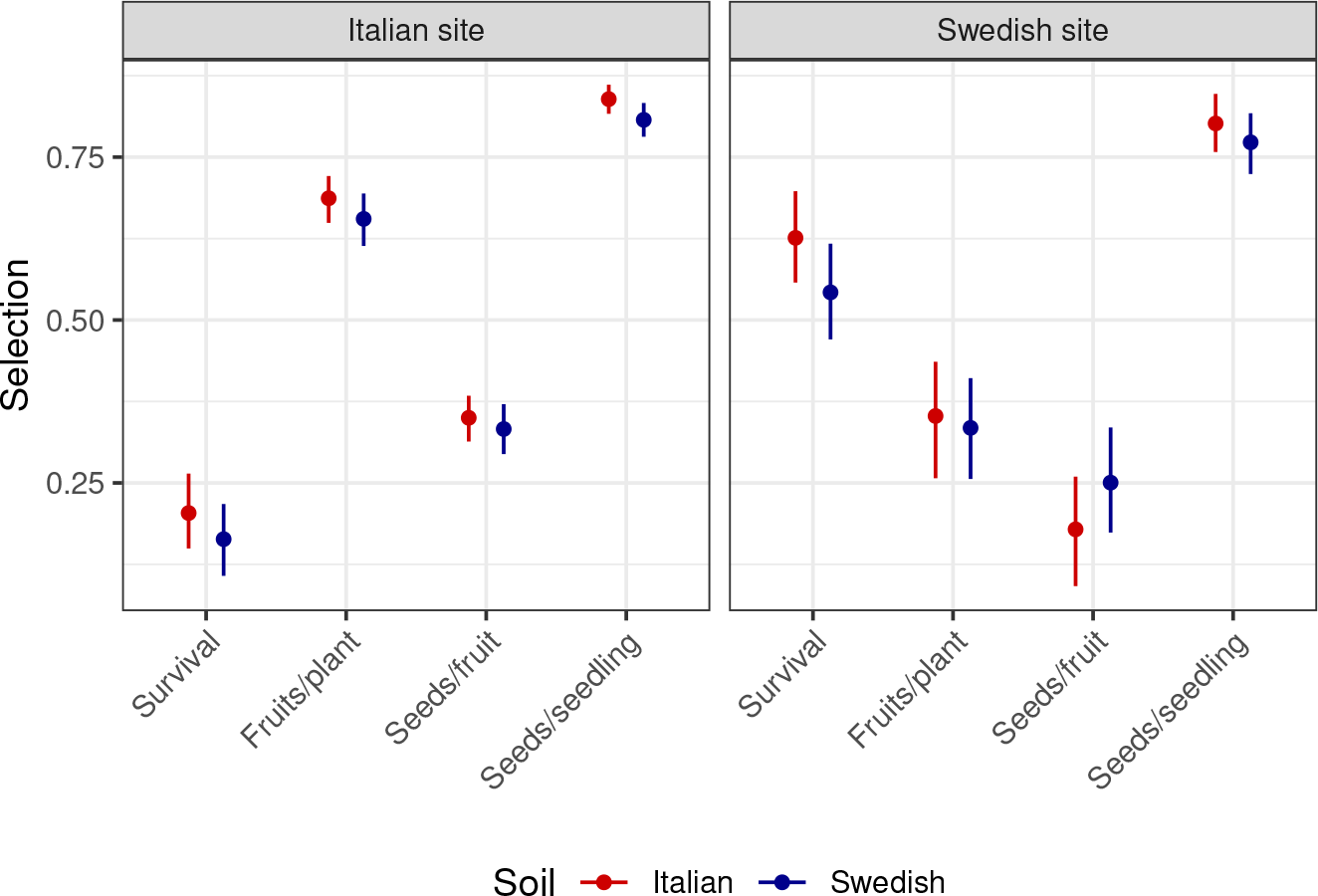
Selection favouring the local ecotype on Italian and Swedish soils at the Swedish and Italian sites. Plots show selection coefficients with 95% confidence intervals for survival to reproduction, number of fruits per reproductive plant, number of seeds per fruit, and number of seeds per planted seedling.

We found only small differences in the strength of selection on the two soil types (fig. 2). At the Italian site selection, selection quantified based on overall fitness was slightly stronger on the local soil (Δ*s*=0.032), and this difference approached, but did not exceed, the threshold for statistical significance (p=0.054). Similarly, estimates of selection based on each of the three fitness components were slightly larger on the local soil than on the non-local soil, but these differences were not statistically significant (p≥0.234). At the Swedish site, the estimate of selection through number of seeds per fruit was slightly larger on the local than on the non-local soil (Δ*s*=0.071), but the opposite was true for estimates of selection through survival, number of fruits per reproductive plant and number of seeds per planted seedling (Δ*s*=0.084, 0.019, 0.029 respectively), although none of these differences were statistically significant (p≥0.088). These results indicate that there were no large differences in the strength of selection between soil types at either site.

## Discussion

This study quantified the contribution of adaptation to soil type to overall adaptive differentiation between two locally adapted populations of *A. thaliana* occurring on contrasting soils and close to the northern and southern range limit in Europe. Although overall fitness differences between local and non-local ecotypes were very strong at both sites, differences in the strength of selection when measured on local and non-local soil types were small and not convincingly different from zero. These observations suggest that adaptation to local soil type makes at most a very limited contribution to local adaptation in these populations.

The results are unlikely to be an artefact of low statistical power. Sample sizes for each site-soil-ecotype treatment were large, and the confidence intervals around estimates of fitness and selection coefficients were narrow (fig. 1, 2). It is plausible that an even larger experiment could have detected a statistically significant difference in the strength of selection between soil types, especially for overall fitness in Italy. However, the absolute magnitude of the difference would remain very small, and as such the biological significance of the result would remain slight/ The main conclusion that adaptation to soil type contributes little to local adaptation would thus remain unchanged.

The absence of evidence for adaptation to local soil contrasts to reports of adaptive differences between *A. thaliana* ecotypes from calcareous and non-calcareous soils (12) and from soils with high and low salt content in Spain (24; 25).

There are at least two possible explanations for the contrasting results. First, the soil types considered in the present study may from the plant’s perspective have been too similar to elicit different responses to soil-mediated selection. Although these soils differ in their chemical composition (13), they are both low in nutrients, and prone to desiccation, a situation likely common to soils in most *A. thaliana* natural habitats. Second, previous work has shown that at the sites of the present study, the strength of selection against the non-local ecotype can vary strongly among years (15; 26), and repeating this experiment in other years may have yielded different results. However, the present results are consistent with those of a reciprocal transplant experiment conducted in the preceding year, which was characterised by markedly stronger selection against the nonlocal ecotype at both sites (13). Taken together, these studies suggest that differential adaptation to soil conditions is at most weak between this pair of ecotypes, reflecting a lack of genetic variation for adaptation to the local soil types studied.

Previous work on the study populations suggest that divergent biotic selection makes a minor contribution to local adaptation between the two study populations, whereas there is strong evidence for differences in climatic conditions playing a major role. A minor role for divergent biotic selection is suggested by the fact that damage by herbivores and pathogens is limited at the native sites of the two populations (J. Ågren and T. Ellis, personal observation), and that no effect of the soil microbiome on the relative fitness of the two ecotypes were detected under controlled conditions (20). By contrast, differences between the two populations in seed dormancy, and phenology of seed germination and flowering are consistent with documented divergent selection on these traits, and with differences between the native sites in the timing of conditions favourable for seedling establishment, growth, and seed production (17; 27; 28). Moreover, in Sweden, minimum winter temperature is a strong predictor of strength of selection against the nonlocal ecotype (15), and local plants show higher freezing tolerance and ability to photosynthesise at low temperatures compared to the Italian ecotype (18; 19; 26). Thus, adaptation to local climatic conditions appears to be the primary driver of adaptive differentiation between these ecotypes. Differences between the two ecotypes in seed dormancy, flowering time, and cold tolerance are consistent with large-scale latitudinal variation in these traits across the European range of *A. thaliana* (29–33). However, additional field experiments will be required to determine to what extent variation in these traits reflects divergent selection among local populations, gene flow, and historical factors across a variety of spatial scales.

## Data and code availability

Raw data, along with full scripts to reproduce the analysis, figures and manuscript are available on GitHub (https://github.com/ellisztamas/soil_adaptation_ellis_agren) and Zenodo (https://doi.org/10.5281/zenodo.11061217).

## Acknowledgments

This work was made possible by the help of M. Vass, L. Vikström, B. Ellis, L. Dück and M. Vinyeta. We also thank P. Falzini and Y. Jonsson for permission to conduct experiments on their land, and to the Botanical Garden of Rome for generously allowing us to use their greenhouse facilities. The study was financially supported by grants from the Swedish Research Council to JÅ (2016-05435, 349-2007-8731).

## Author contributions

TJE performed the experiment and analysis, and wrote the manuscript. JÅ conceived the study, and critically revised the manuscript.

## Competing interests

The authors declare no competing interests.

## Notes

https://doi.org/10.5281/zenodo.11061217

### Competing Interest Statement

The authors have declared no competing interest.

### Summary of Updates

The updated version includes Swedish-research-council grant numbers awarded to Jon Agren.

